# Spared nerve injury causes motor phenotypes unrelated to pain in mice

**DOI:** 10.1101/2023.07.07.548155

**Authors:** Makenzie R. Norris, Samantha S. Dunn, Bhooma R. Aravamuthan, Jordan G. McCall

**Author notes:** To whom correspondence should be addressed: B.R.A.: Address: 660 S. Euclid Ave, Box 8111, St. Louis, MO, USA, 63110. Website: www.aravamuthanlab.wustl.edu, J.G.M.: Address: 660 S. Euclid Ave, Box 8054, St. Louis, MO, USA, 63110. Website: www.mccall-lab.org.

## Abstract

Most animal models of neuropathic pain use targeted nerve injuries quantified with motor reflexive measures in response to an applied noxious stimulus. These motor reflexive measures can only accurately represent a pain response if motor function in also intact. The commonly used spared nerve injury (SNI) model, however, damages the tibial and common peroneal nerves that should result in motor phenotypes (i.e., an immobile or “flail” foot) not typically captured in sensory assays. To test the extent of these issues, we used DeepLabCut, a deep learning-based markerless pose estimation tool to quantify spontaneous limb position in C57BL/6J mice during tail suspension following either SNI or sham surgery. Using this granular detail, we identified the expected flail foot-like impairment, but we also found SNI mice hold their injured limb closer to the body midline compared to shams. These phenotypes were not present in the Complete Freunds Adjuvant model of inflammatory pain and were not reversed by multiple analgesics with different mechanisms of action, suggesting these SNI-specific phenotypes are not directly related to pain. Together these results suggest SNI causes previously undescribed phenotypes unrelated to altered sensation that are likely underappreciated while interpreting preclinical pain research outcomes.

## Introduction

Preclinical pain studies have not always translated into efficacious human therapeutics^1^. Considering this shortfall, it is critical that we deeply evaluate the validity of common neuropathic pain models to improve our interpretations of these studies and their implications. The spared nerve injury (SNI) model is widely used to model neuropathic pain in rodents due to its resulting robust and prolonged mechanical hypersensitivity^2–10^. Despite its common use, however, SNI has several limitations including gait changes that may not be pain-related and potential lack of direct clinical validity^8, 11^. It is additionally notable that the SNI injury (tibial and personal nerve ligation) should clinically cause a flail-foot deformity, or absence of the ability to dorsiflex or plantarflex at the ankle. This motor symptom, if seen in the SNI model, would significantly limit the ability to interpret some sensory assays (e.g., voluntary hindpaw withdrawal assays). Here we use DeepLabCut, a deep learning tool used to mark body position and tracking movements to quantify variability in limb movement of injured and non-injured hindlimbs^12^. We readily identified a flail foot-like presentation, but also observed an unexpected postural phenotype we subsequently found to be unrelated to both flail foot and mechanical hypersensitivity. These newly observed phenotypes may affect the response to sensory assays are important to identify and distinguish from pain responses to ensure the ongoing value of the SNI model for assessing neuropathic pain in mice.

## Materials and Methods

### Animals

All experiments and procedures were approved by the Animal Care and Use Committee of Washington University School of Medicine in accordance with National Institutes of Health guidelines. Adult male and female C57BL/6J mice were used from age 8-15 weeks. All mice were group-housed, given *ad libitum* access to standard laboratory chow (PicoLab Rodent Diet 20, LabDiet, St. Louis, MO, USA) and water, and maintained on a 12:12-hour light/dark cycle (lights on at 7:00 AM). Experimenters were blinded to mouse conditions including sex and injury status during experimental data collection and analysis.

### Spared Nerve Injury (SNI)

The surgical procedure for the SNI-induced model of neuropathic pain was performed as described previously^4, 10^. Mice were anesthetized with 3% isoflurane and right hind limb shaved and disinfected with 75% ethanol and betadine. A 10-15 mm incision was made in the skin proximal to the knee to expose and separate the biceps femoris muscle. The common peroneal and tibial branches were ligated with 6-0 silk suture (Ethicon Inc., Raritan, NJ, USA) and 1 mm of nerve was excised distal to the ligature, leaving the sural branch intact. Following wound closure mice recovered on a 43°C table prior to returning to their home cage. Sham surgeries were identical to the SNI procedure without the ligation, excision, and severing of the peroneal and tibial branches. Behavioral testing began on post-operative day 7 and wound clips were removed from the healed incision after testing was completed on post-operative day 7.

### Complete Freund’s Adjuvant (CFA)

Mice were intraplantarly injected with resuspended 100 µl of CFA (Thermo Fischer Scientific, Waltham, MA, USA) or 0.9% saline with a 26 gauged needle coupled to a 1 mL syringe as described previously^13^. Behavioral testing on these animals began on post-injection day 1.

### Mechanical sensitivity (von Frey)

Mice were acclimated for 2 hours on an elevated wire mesh grid in 5-inch diameter plexiglass cylinders wrapped in black opaque plastic sheets. Mechanical hypersensitivity was determined by applying von Frey filaments (0.02 g to 3.5 g; Bioseb, Pinellas Park, FL,USA) to the lateral aspect of the hindpaw^14^ using the up-down method and 50% withdrawal threshold was calculated as described previously^10, 15^.

### Tail Suspension

Mice were suspended by their tail for 1 minute with a clear plastic cylinder placed at the tail base to avoid tail climbing behavior^16^. Videos of tail suspension were taken at 120 frames per second using Google Pixel 3 (Google, Mountain View, CA, USA) smartphones. Tail suspension was performed immediately after von Frey testing on days without pharmacological interventions. When pharmacological interventions were used, von Frey and tail suspension data were taken on separate days; tail suspension first with at least two days of drug/vehicle washout in between treatments.

### Pose estimation during tail suspension

DeepLabCut (version 2.1.1) was used to track the following points for each mouse in two-dimensional space: tailbase, trunk midpoint at the plastic cylinder base, right and left hindpaw heel, and right and left hindpaw middle toes. Using k-means-based extraction, 20 frames were extracted from a subset of 56 tail suspension videos taken from different experimental days, experimental cohorts, and sexes. The above points were manually labeled on these 1,120 frames and then used to train a ResNet-50-based neural network for 1,030,000 training iterations to achieve a train error less than 3 pixels and a test error less than 8 pixels (frame size 1920×1080 pixels, ∼7 pixels/mm). The trained model was assessed for accuracy by visually checking 3 videos. A p-cutoff of 0.9 was used to condition the X,Y coordinates of each video for further analysis.

Median foot movement and distance from the mouse trunk midline were calculated using the above tracked two-dimensional coordinates and code custom written in MATLAB (R2021, The MathWorks, Inc). These metrics were: median distance moved of the right and left hindpaw heel relative to the mouse tailbase over the course of the 1 minute tail suspension video, median distance moved of the right and left hindpaw middle toe relative to the mouse tailbase, median distance moved of the right and left hindpaw toe relative to the ipsilateral hindpaw heel, median distance of the right and left hindpaw heels to the mouse midline (distance perpendicular to a line drawn between the mouse tailbase and trunk midpoint), and median distance of right and left hindpaw toes to mouse midline. Given natural variation in mouse size, especially by sex, these distances are all presented as ratios relative to foot length.

### Drugs

Fenobam (30 mg/kg, i.p.; HelloBio, Princeton, NJ, USA) was dissolved in 100% dimethyl sulfoxide on the day of experiment in volumes of 20 µl^7^. Behavioral experiments were conducted 5 minutes post-Fenobam injection. Gabapentin (30 mg/kg, i.p.; Sigma-Aldrich, St. Louis, MO, USA) was dissolved in 0.9% saline on the day of experiment. Gabapentin was injected i.p. 1 hour prior to behavioral testing^8^. Metformin (LKT Laboratories, St. Paul, MN, USA) and dissolved in 0.9% saline freshly for each use. Metformin was injected at 200 mg/kg daily for 7 consecutive days between 10am-1pm^5^. Behavioral experiments were conducted at least 24 hours after the last metformin injection. Carprofen (Zoetis, Kalamazoo, MI, USA) was administered i.p.,1 hour prior to behavioral testing at 50 mg/kg.

### Statistical Analysis

All data are expressed as mean ± SEM. Differences between normally distributed groups were determined using mixed-effects two-way ANOVA analysis followed by posthoc Sidak’s comparison when the main effect was significant (p < 0.05). Statistical analyses were conducted using Prism 8.2 (GraphPad).

#### Data availability

All data presented in this manuscript is available in **Supplemental Table 1**

## Results

### Objective quantification using deep learning following SNI induced long-term mechanical hypersensitivity

The SNI model of neuropathic pain has been shown to reliably induce ipsilateral mechanical hypersensitivity^2–10^. Here, we performed SNI or sham surgeries and used the von Frey assay to quantify mechanical withdrawal thresholds (Fig. 1A&B). As expected, SNI induced ipsilateral hindpaw mechanical hypersensitivity. Similar assessment of the contralateral hindpaw showed a gradual decrease in mechanical withdrawal threshold in SNI mice during week four that returned to baseline by week six. (Fig. 1C). Each week, we briefly suspended the animals by the tail to enable quantification of hind limb positions using DeepLabCut. To quantify postural changes in mice, we suspended mice by their tail and recorded the ventral portion of the body for 1 minute at 120 frames/sec (Fig. 1D). K-means extraction was used to extract a subset of frames (Fig. 1E) which were then manually labelled for body parts of interest for each hindlimb (Fig. 1F). From these labelled frames, we trained a neural network to identify body postural points (Fig. 1G). Lastly, we used spatial coordinates provided by our previously trained neural network to calculate limb position metrics across all frames (Fig. 1H).

**Figure 1:**
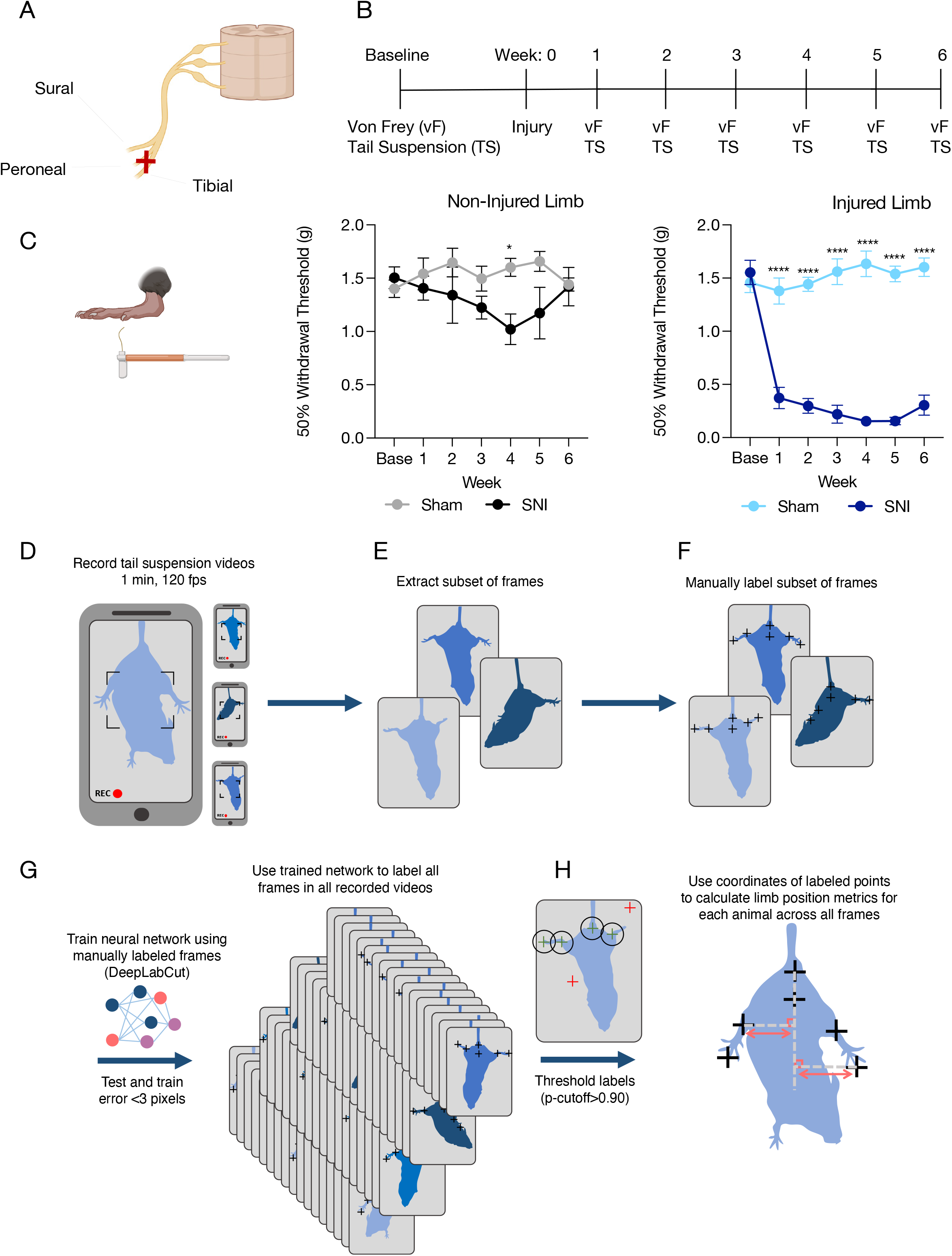
DeepLabCut-based approach to quantifying limb position following SNI-induced mechanical hypersensitivity. (**A**) Cartoon depicting SNI approach. (**B**) Calendar showing experimental timeline. (**C**) SNI significantly decreases 50% withdrawal threshold throughout six-week timecourse. Data represented as mean +/−SEM; n=18-20/group; two-way ANOVA, Sidak’s post hoc (non-injured limb Sham versus SNI; Injured limb Sham versus SNI *p<0.05 ****p<0.0001). **(D)** Recording of tail suspension for 1 min at 120 frames per second (fps). **(E)** K-means-based extraction of a subset of frames. **(F)** Manual labeling of the tailbase, trunk midpoint, right and left hindpaw heel, and right and left hindpaw middle toes on the extracted frames. **(G)** Manually labeled frames are used to train a ResNet-50-based neural network to label the remaining frames. **(H)** A threshold p-cutoff of 0.9 was used to condition the X,Y coordinates of each video for calculation of median foot movement and distance from the mouse trunk midline.

### SNI decreases contralateral limb movement and causes flail foot-like phenotype

To determine whether SNI causes the predicted flail foot phenomenon, we focused on toe and heel movement changes relative to body midline and toe movement relative to the heel. Quantifying toe movement relative to the body midline showed that SNI decreased overall toe and heel movement in the contralateral, but not ipsilateral hindlimb for the duration of the injury (Supplementary Fig. 1A-F), perhaps suggesting a compensatory motor coordination phenomenon in the uninjured limb following SNI. As would be expected with flail foot (decreased movement at the ankle), SNI decreased toe movement relative to the heel in the ipsilateral injured hindlimb but also in the contralateral hindlimb (Fig. 2A-C) again suggesting compensatory SNI-induced movement changes in the contralateral limb. This decreased toe movement emerges within a week of SNI, but unlike the mechanical hypersensitivity (Fig. 1C), appears to resolve after six weeks. Given the presence of these previously uncharacterized postural phenotypes, we next assessed for other latent postural phenotypes in the SNI model.

**Figure 2:**
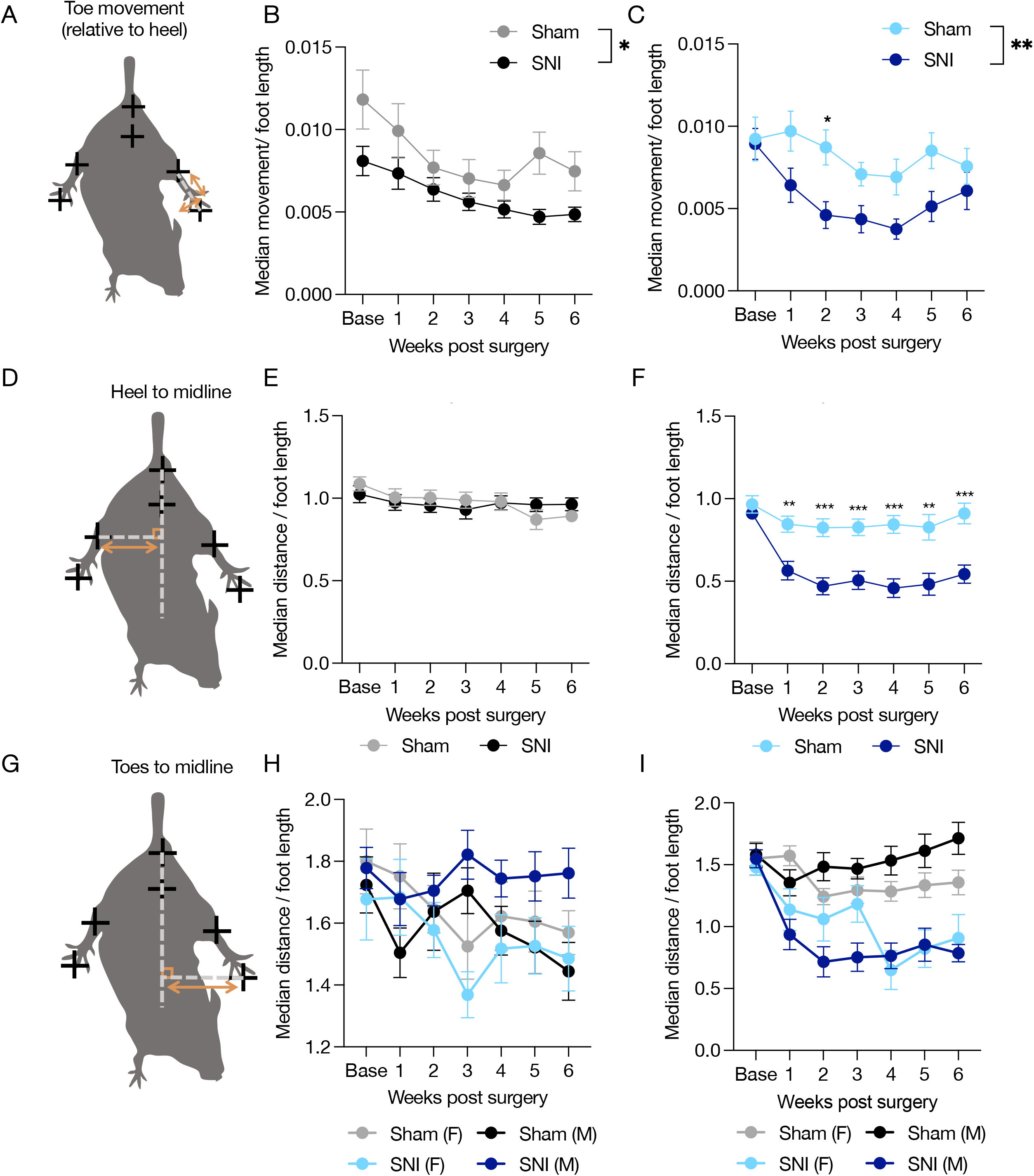
SNI induces flail foot in injured limb and causes injured limb to be held closer to body midline. (**A**) Image shows points of comparison quantified in (**B**) and (**C**). (**B**) and (**C**) show median movement between the toe of non-injured (**B**) and injured (**C**) limbs relative to the heel. (**C**) shows a significant main effect between Sham and SNI on injured limb side, a phenomenon we call “flail” foot. Data from all comparisons show a significant decrease in movement on non-injured sides. All data expressed as mean +/−SEM; n=18-20/group; two-way ANOVA, Sidak’s post hoc (Non-injured limb Sham versus SNI; Injured limb Sham versus SNI **p<0.01 ***p<0.001, ****p<0.0001). (**D**) Image shows points of comparison quantified in (**E**) and (**F**). (**E**) and (**F**) show median distance between heel of non-injured (**E**) and injured (**F**) limb and midline. SNI significantly reduces distance between heel and body midline during tail suspension. Data expressed as mean +/−SEM; n=18-20/group; two-way ANOVA, Sidak’s post hoc (Non-injured limb Sham versus SNI; Injured limb Sham versus SNI **p<0.01 ***p<0.001, ****p<0.0001). (**G**) Image shows points of comparison quantified in (**H**) and (**I**). (**H**) and **(I**) show median distance between toe of non-injured and injured limb and midline divided by sex. (**H**) shows a three-way significant interaction between Week/Pain/Sex suggesting the time course of SNI-induced decreased toes to midline distance is significantly different between males and females. Data represented as mean +/−SEM; n= 8-12/group; three-way ANOVA (Week x Pain x Sex), Tukey post hoc (Pain vs. Sex vs. Time *p<0.05).

### SNI decreases the distance between the injured limb and body midline

To determine whether SNI causes other postural changes, we next quantified the distance between heel and midline in sham and SNI mice (Fig. 2D). We found SNI did not significantly change heel to midline distance on the contralateral limb (Fig. 2E). However, we found a significant decrease in distance between heel and midline on the ipsilateral injured limb across all six weeks of tail suspension testing (Fig. 2F). Similarly, we found no significant effect of SNI on contralateral toe position relative to midline (Fig. 2G&H). However, SNI decreased the distance of the ipsilateral toes to midline in a sex-dependent manner. Specifically, male mice significantly decreased the distance between toes and body midline across all six weeks, but female mice did not develop this same decrease in distance between toes and body midline until week four of tail suspension testing (Fig. 2I). This suggests the temporal impact of SNI differs between males and females, in line with what others have seen in other behavioral paradigms^6, 17–19^. Importantly, this was the only sex difference we found throughout this entire study. We next used 15 second bins to determine if the full duration of the recording was necessary. Here, while the distance from midline is not affected during baseline, animals do hold their limbs closer to midline in the first 15 seconds at week one and all subsequent bins through the following weeks (Supplementary Fig. 2). Together, SNI stably decreases overall distance between the injured limb and body midline for at least six weeks following injury.

### Acute inflammatory injury does not decrease distance between injured limb position and body midline

Upon manual observation of the DeepLabCut-tracked video files, we noted that many SNI animals appeared to be performing a guarding-like behavior (Supplementary Fig. 3). To determine whether this limb position reflected guarding, we used the Complete Freund’s Adjuvant (CFA) model of inflammatory hindpaw pain known to cause guarding in rodents^20^. We reasoned that this approach would allow us to simultaneously determine the selectivity of this phenotype for SNI compared to a completely different pain model as well as provide insight into the relationship to guarding. CFA decreased mechanical withdrawal threshold using the Von Frey assay (Fig. 3A-C) which was reversed using the non-steroidal anti-inflammatory carprofen (Fig. 3C). To determine whether SNI-induced changes to limb position were specific to that injury, we quantified the distance between heel and midline in mice with an acute inflammatory injury. Unlike SNI, CFA did not alter heel to midline distance or toe position for either the non-injured or injured limbs (Fig. 3D-I). This suggests the decrease in injured limb position relative to body midline effect we show in Fig. 2 is specific to SNI and does not reflect a broad form of injury guarding.

**Figure 3:**
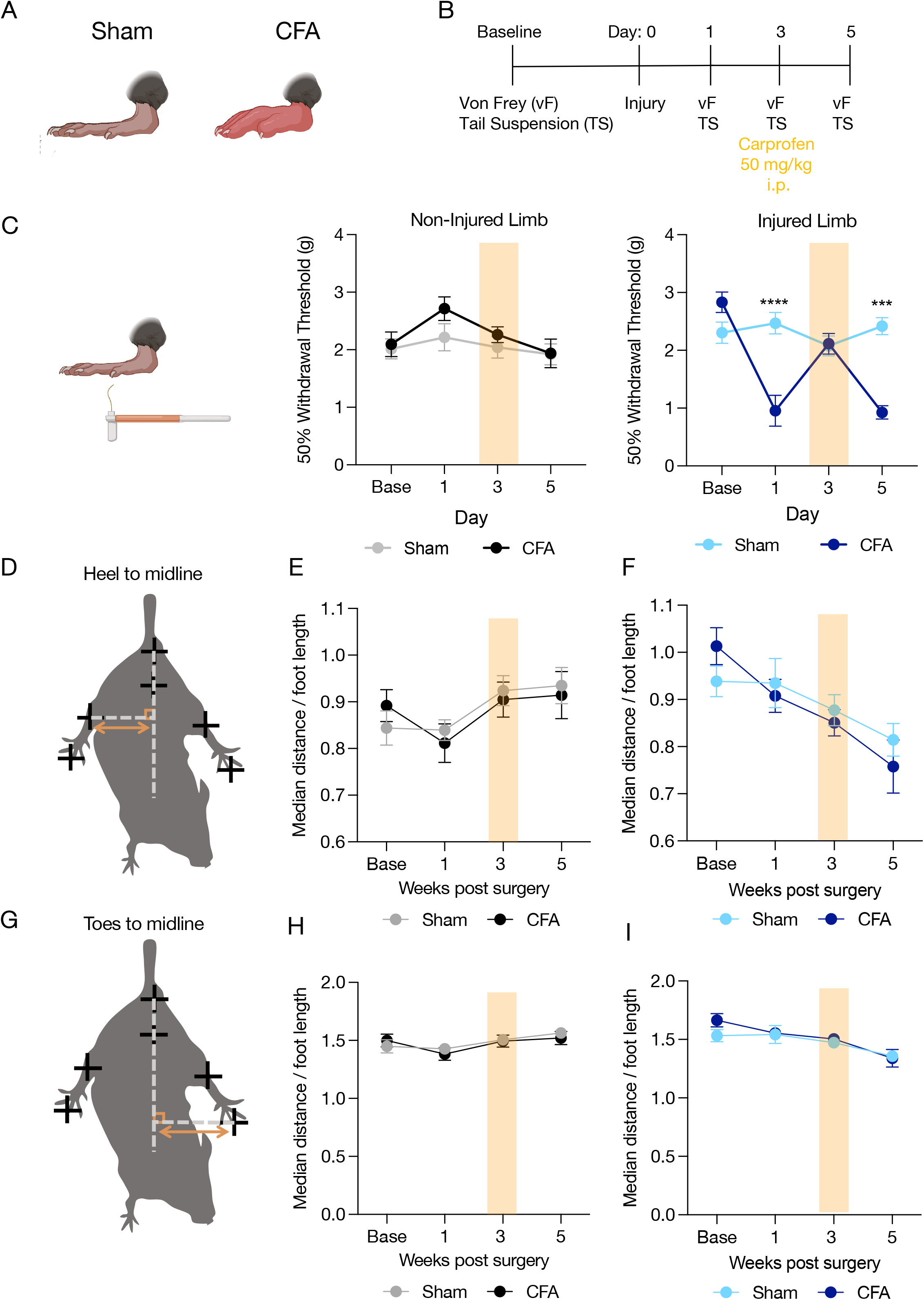
Acute inflammatory injury does not decrease distance between injured limb and body midline. (**A**) Cartoon shows diagram of inflammatory injury. (**B**) Calendar shows experimental timeline and analgesic drug dosages used. (**C**) High-dose carprofen reverses CFA-induced mechanical hypersensitivity. data expressed as mean +/−SEM; n=20/group; two-way ANOVA, Sidak’s post hoc (Non-injured limb Sham versus CFA; Injured limb Sham versus CFA ***p<0.001, ****p<0.0001). (**D**) Images shows points of comparison quantified in (**E**) and (**F**). (**G**) Image shows points of comparison quantified in (**H**) and (**I**). CFA did not decrease distance between toes/heel to midline as shown in SNI. Data represented as mean +/−SEM; n=20/group; two-way ANOVA, Sidak’s post hoc (Non-injured limb Sham versus CFA; Injured limb Sham versus CFA.

### Analgesic treatment does not prevent SNI-induced decreased distance between injured limb and body midline

If this SNI-decreased distance between injured limb and body midline is a pain-related phenotype an analgesic should reverse its course. To test this hypothesis, we administered several clinically-used or preclinically tested analgesics known to reverse SNI-induced mechanical hypersensitivity^5, 7, 8^. We first chose the anticonvulsant gabapentin as it has previously reversed both SNI-induced mechanical hypersensitivity as well as gait-related phenotypes (Supplementary Fig. 4A-C)^8^. Gabapentin (30 mg/kg i.p.) did not blunt the SNI-induced decreased distance between the injured limb and body (Supplementary Fig. 4D-I), but significantly decrease movement across all measures and limbs during gabapentin treatment (Supplementary Fig. 5) suggesting potential gabapentin-induced sedation. In the next cohort we tested different analgesics with differential mechanisms of action including the metabotropic glutamate receptor negative allosteric modulator, fenobam (30 mg/mg i.p.)^7^, the AMP-activated protein kinase activator metformin (seven consecutive days of 200 mg/kg i.p.)^5^, and we re-tested gabapentin interleaved with vehicle treatments (Fig. 4A). We found fenobam, gabapentin, and metformin administration all had no effect on mechanical thresholds for the non-injured limb, but each reversed SNI-induced mechanical hypersensitivity in the injured hind limb (Fig. 4B). Despite this reversal, none of the analgesic treatments altered non-injured or injured limb position (Fig. 4C-H) or the relative movements of this body parts representing the flail foot-like observation (Supplementary Fig. 6A-I). Together, this data suggests the newly observed decrease in heel to midline distance likely does not represent a pain-related behavioral phenotype.

**Figure 4:**
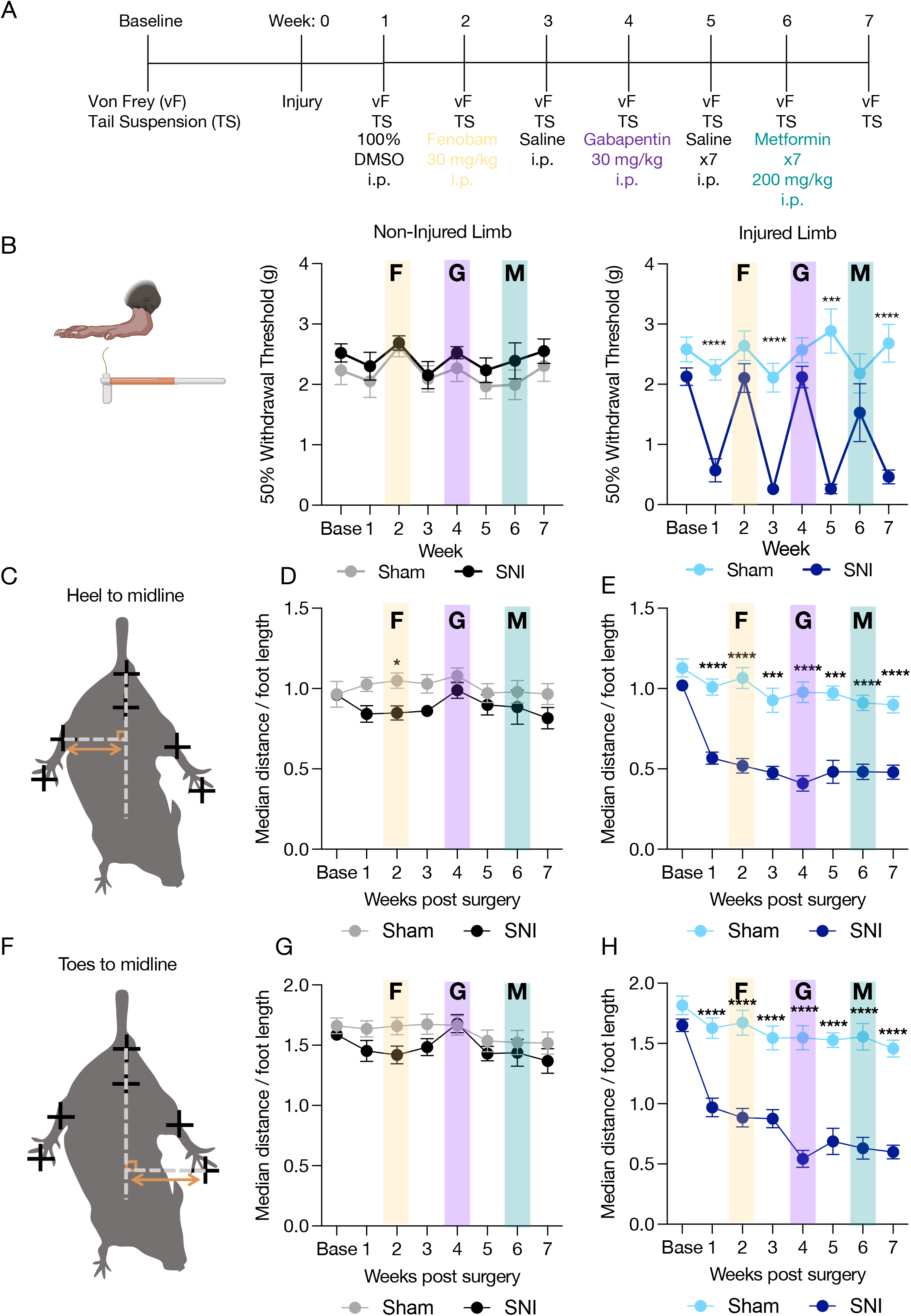
Analgesics do not reverse SNI-induced decrease in limb distance from body midline. (**A**) Calendar shows experimental timeline and analgesic drug dosages used. (**B**) Fenobam, Gabapentin, and Metformin all reverse SNI-induced mechanical hypersensitivity (**C**) Images shows points of comparison quantified in (**D)** and (**E**). Analgesics did not reverse SNI-induced decrease in heel to midline distance, shown in (**D**) and (**E**). (**F**) Image shows points of comparison quantified in (**G**) and (**H**). Analgesics did not reverse SNI-induced decrease in toes to midline distance, shown in (**G**) and (**H**). Data represented as mean +/−SEM; n=15-21/group; two-way ANOVA, Sidak’s post hoc (Non-injured limb Sham versus SNI; Injured limb Sham versus SNI *p<0.05 ***p<0.001 ****p<0.0001). Metformin data does not include female mice as its analgesic efficacy is limited to males (n=10-12/group). F = Fenobam, G = Gabapentin, M= Metformin.

## Discussion

We conducted an in-depth investigation into SNI-associated postures that identified a flail foot-like phenotype consistent with human tibial/peroneal paralysis as well as an unexpected decrease in injured limb distance from body midline. These neuropathic injury-specific postural changes help guide the interpretation of preclinical pain studies. One recent study has identified proprioceptive deficits in Nav1.1 null mice that resulted in postural phenotype like the changes identified in SNI mice in our study^21^. Voltage-gated sodium channels play a critical role in the transmission of pain signals and Nav1.1 has been associated with paroxysmal extreme pain disorder (PEPD)^22^. Future work should investigate the association between Nav1.1 and the neuropathic pain-specific postural changes shown here. This study was performed entirely with postural information during tail suspension, further identification of potential proprioceptive abnormalities in freely moving SNI mice will help extend our understanding of these implications. Previously these types of analyses have been hindered by a lack of reliable tools. However, recent advances in animal tracking will likely aid in overcoming this hurdle^23–30^. Furthermore, while relevant to the injury, the change in position of the animal could be exacerbated by acute changes in sympathetic tone to redistribute blood flow due to being held upside down. To determine whether the observed postural changes were specific to neuropathic injury, we used the CFA inflammatory model as a positive control for guarding. These results suggested specificity of postural changes to SNI, but it will be important to compare to other nerve injury models to better differentiate behavioral states specific to each injury. Altogether, this work highlights a need to better understand the nuances of preclinical models of complex human pain syndromes.

## Acknowledgements

We thank the other members of the Al-Hasani and McCall labs for helpful feedback on this project. This work was financially supported by the National Institutes of Health (R01NS117899, J.G.M.; F31NS124301, M.R.N.; K08NS117850, B.R.A), and the Rita Allen Foundation (J.G.M.) with added financial help from the Open Philanthropy Project (J.G.M.). We would like to acknowledge biorender.com for figure cartoons.

## Author contributions

M.R.N, B.R.A, and J.G.M. conceived the project and designed the detailed experimental protocols. M.R.N. and S.S.D. performed the mouse experiments. M.R.N., S.S.D, B.R.A., and J.G.M. performed the investigation and analyzed the data. M.R.N, B.R.A, and J.G.M wrote the paper. M.R.N, S.S.D., B.R.A, and J.G.M edited the paper. M.R.N, B.R.A, and J.G.M. acquired funding. J.G.M. provided research supervision and overall project administration. All authors discussed the results and contributed to revision of the manuscript.

---

**Supplemental Figure 1:**
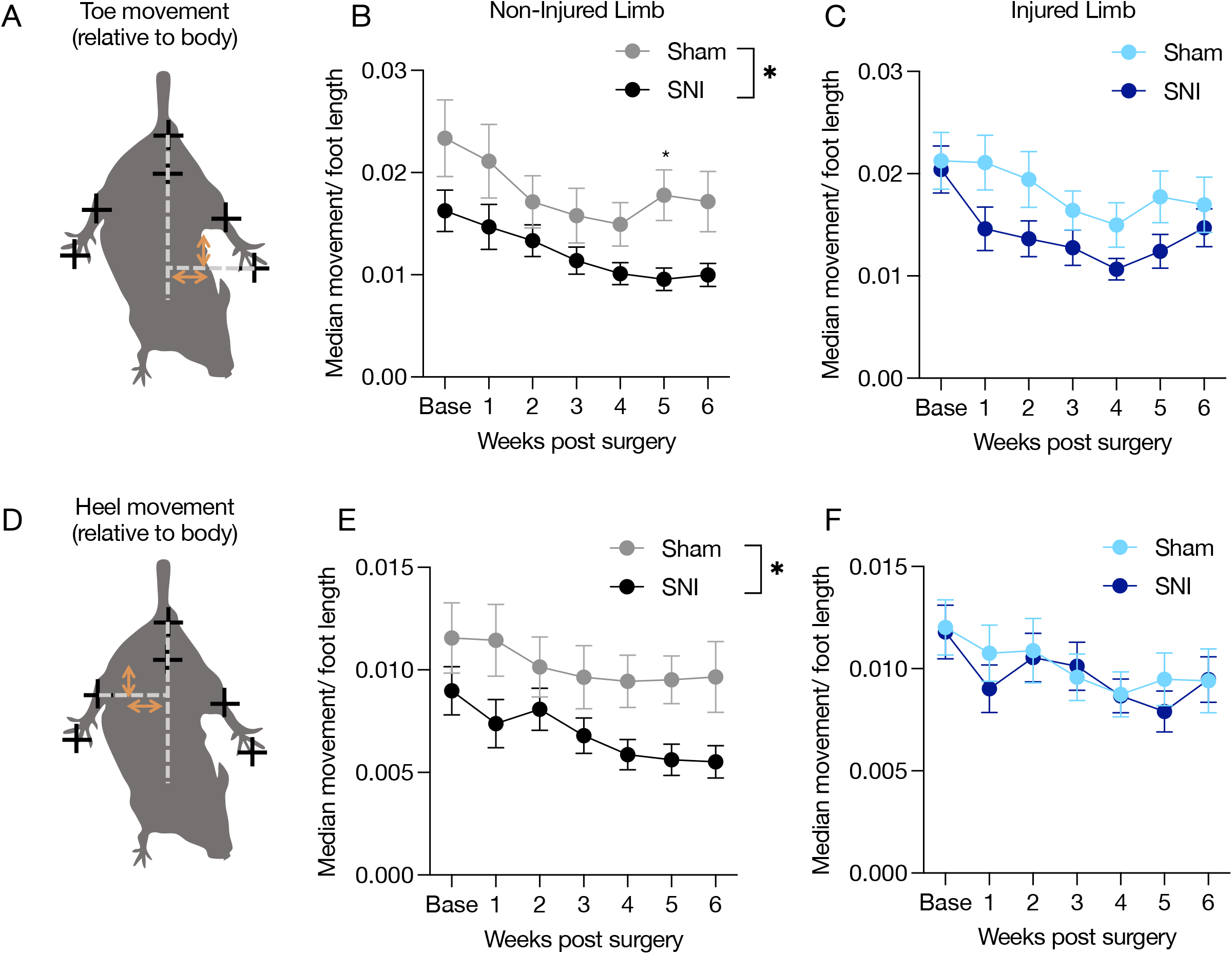
Spared Nerve Injury induces decrease in non-injured limb movement and flail foot in injured limb. (**A**) Image shows points of comparison quantified in (**B**) and (**C**). (**B**) and (**C**) show median movement between the toe of non-injured (**B**) and injured (**C**) limbs relative to body midline. (**D**) Image shows points of comparison quantified in (**E)** and (**F**). (**E**) and (**F**) show median movement between heel of non-injured (**B**) and injured (**C**) limbs relative to midline.

**Supplement Figure 2:**
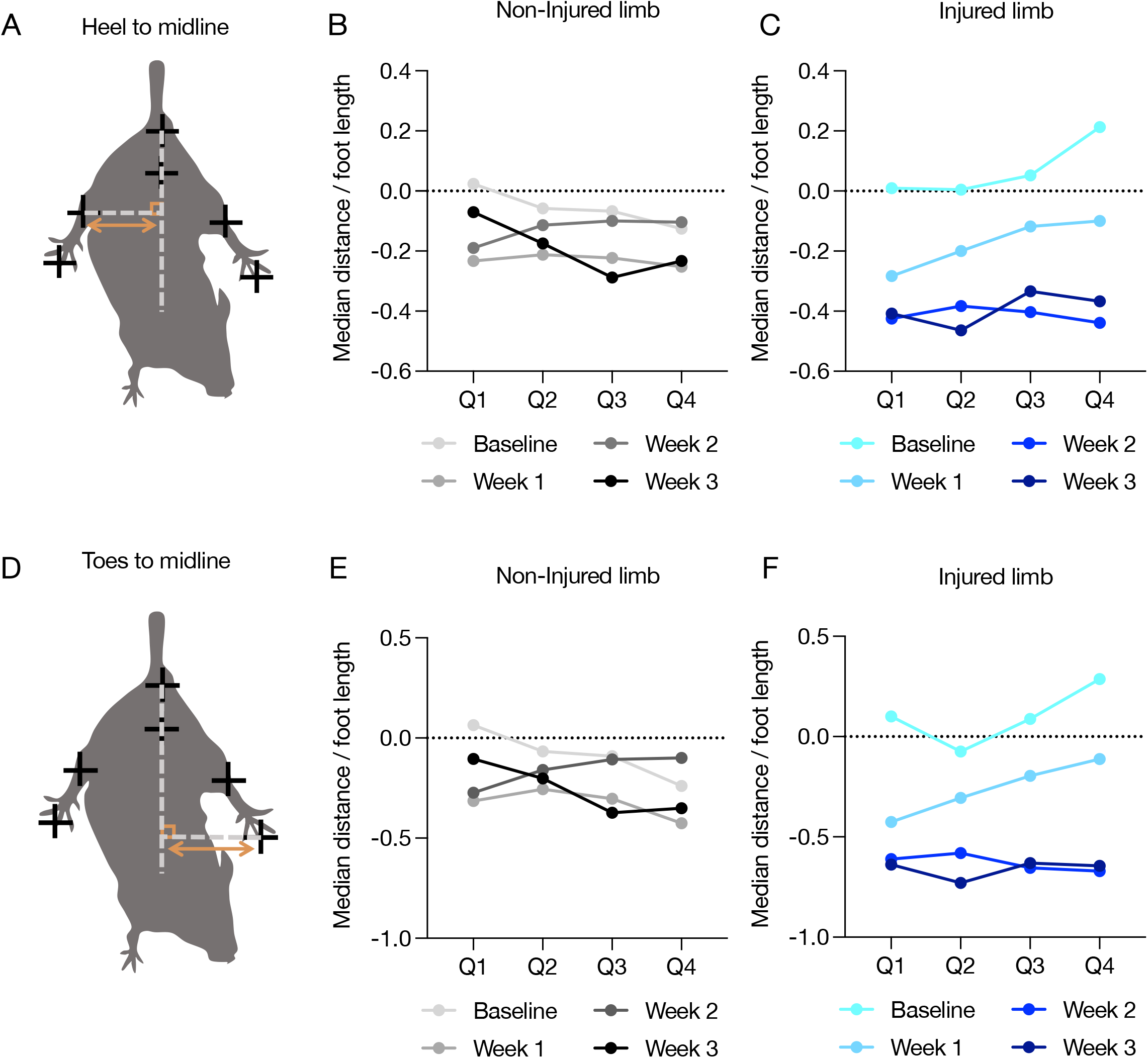
SNI-induced decrease in injured limb distance to body midline is not incremental. (**A**) and (**D**) Image shows points of comparison quantified in (**B**),(**C**) and (**E**),(**F**). (**B**), (**C**), (**E**) and (**F**) separate non-injured (**B**) (**E**) and injured (**C**) (**F**) limbs and show 1 minute tail suspension tests split into four quartiles. Data represented as Sham average – SNI average n=20/group. Week 2 lines correspond to 30 mg/kg i.p. gabapentin administration.

**Supplement Figure 3:**
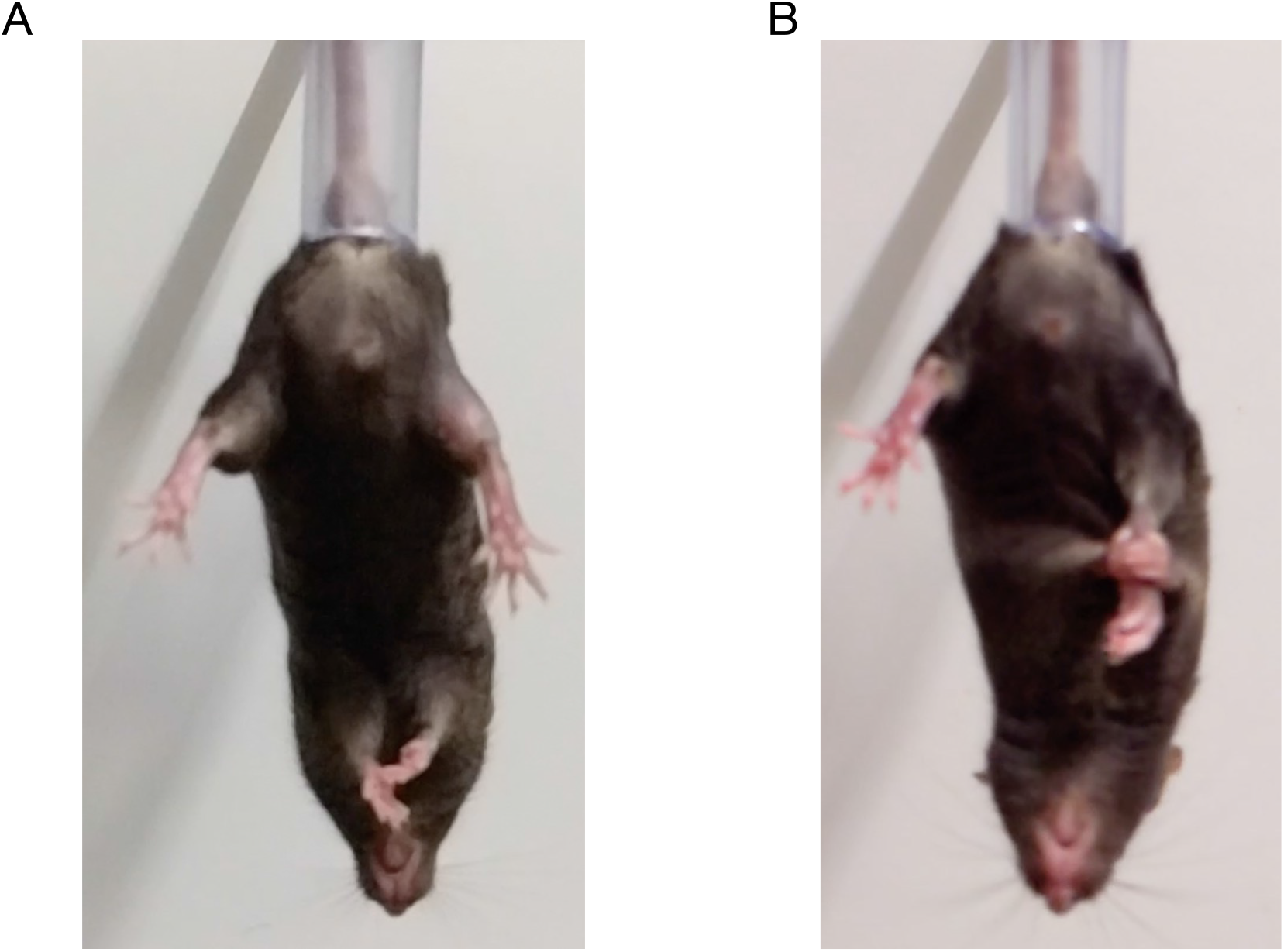
SNI-induced decrease in injured limb distance to body midline resembles guarding. Displays of sham (**A**) and SNI (**B**) animals being suspended by their tails.

**Supplement Figure 4:**
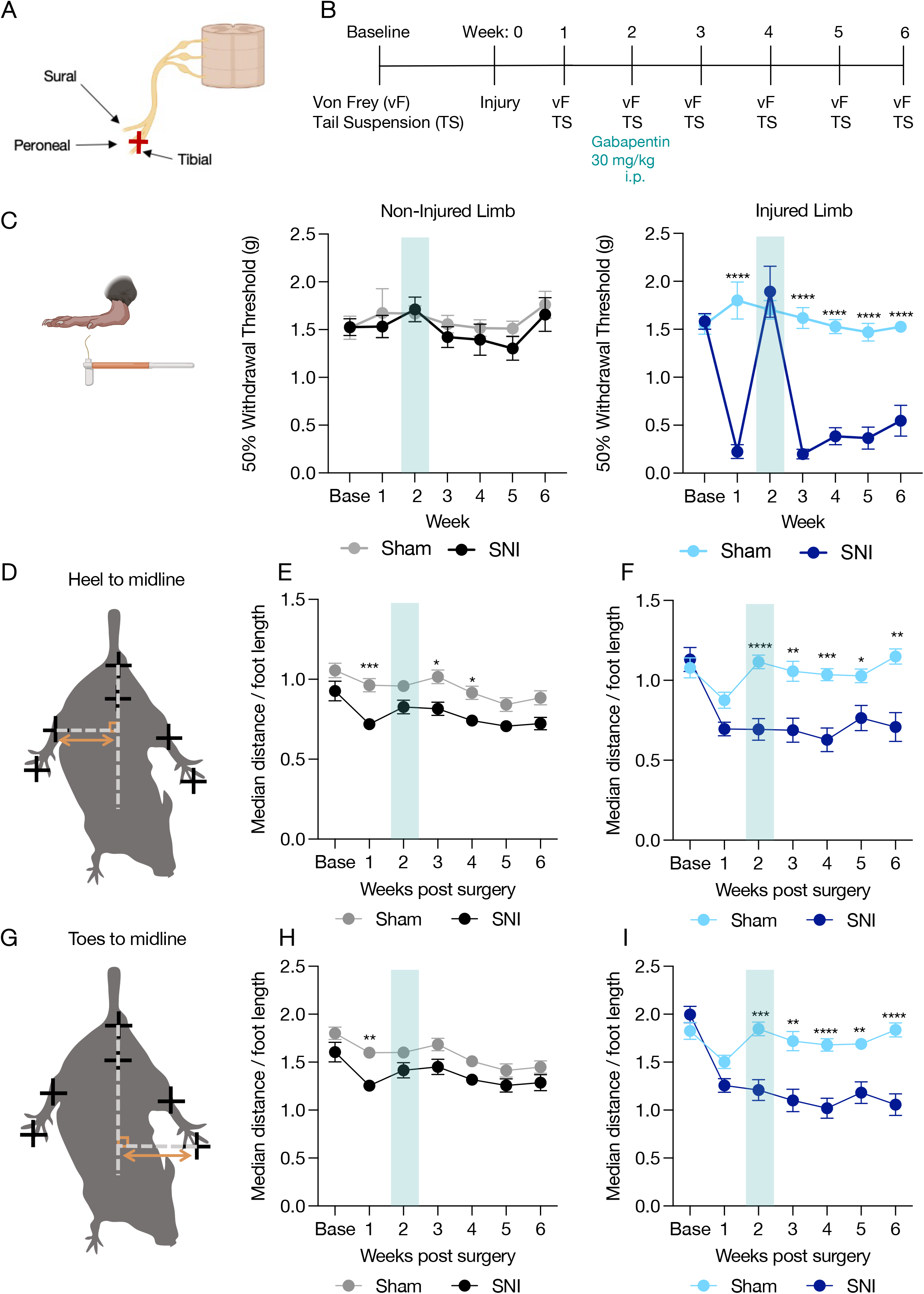
Gabapentin reverses SNI-induced mechanical hypersensitivity but does not reverse SNI-induced decrease in injured limb distance from body midline. (**A**) Cartoon shows diagram of neuropathic injury. (**B**) Calendar shows experimental timeline. (**C**) 30 mg/kg i.p. gabapentin administration reverses SNI-induced mechanical hypersensitivity. Data represented as mean +/−SEM; n=20/group; two-way ANOVA, Sidak’s post hoc (Non-injured limb Sham versus SNI; Injured limb Sham versus SNI ****p<0.0001) (**D**) Image shows points of comparison quantified in (**E**) and (**F**). (**E**) and (**F**) show median distance between heel of non-injured (**E**) and injured (**F**) limb and midline. Spared nerve injury significantly reduces distance between heel and body midline during tail suspension, but this effect is not reversed by gabapentin administration. (**G**) Image shows points of comparison quantified in (**H**) and (**I**). (**H**) and (**I**) show median distance between toe of non-injured (**H**) and injured (**I**) limb and midline divided by sex. Data expressed as mean +/−SEM; n=20/group; two-way ANOVA, Sidak’s post hoc (Non-injured limb Sham versus SNI; Injured limb Sham versus SNI *p<0.05 **p<0.01 ***p<0.001 ****p<0.0001). Toe movement (relative to body)

**Supplement Figure 5:**
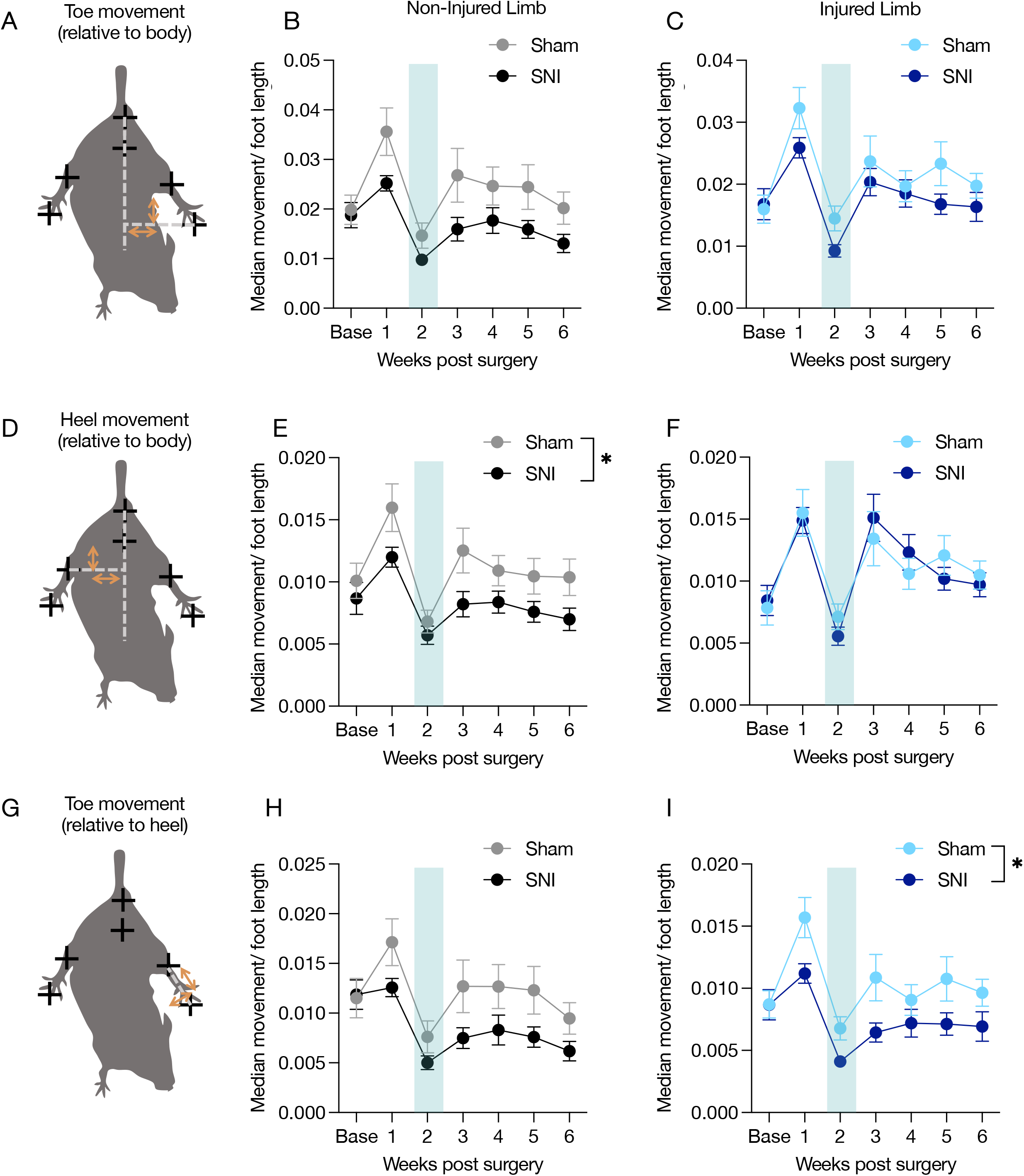
Gabapentin does not reverse SNI-induced decrease in non-injured limb movement and flail foot in injured limb. (**A**) Image shows points of comparison quantified in (**B**) and (**C**). (**B**) and (**C**) show median movement between the toe of non-injured (B) and injured (**C**) limbs relative to body midline. (**D**) Image shows points of comparison quantified in (E) and (**F**). (**E**) and (**F**) show median movement between heel of non-injured (**B**) and injured (**C**) limbs relative to midline. (**G**) Image shows points of comparison quantified in (**H**) and (**I**). (**H**) and (**I**) show median movement between the toe of non-injured (H) and injured (I) limbs relative to the heel. (**I**) shows a significant main effect between Sham and SNI on injured limb side. Data from all comparisons show a significant decrease in movement on non-injured sides. All data expressed as mean +/−SEM; n=20/group; two-way ANOVA, Sidak’s post hoc (Non-injured limb Sham versus SNI; Injured limb Sham versus SNI *p<0.05).

**Supplemental Figure 6:**
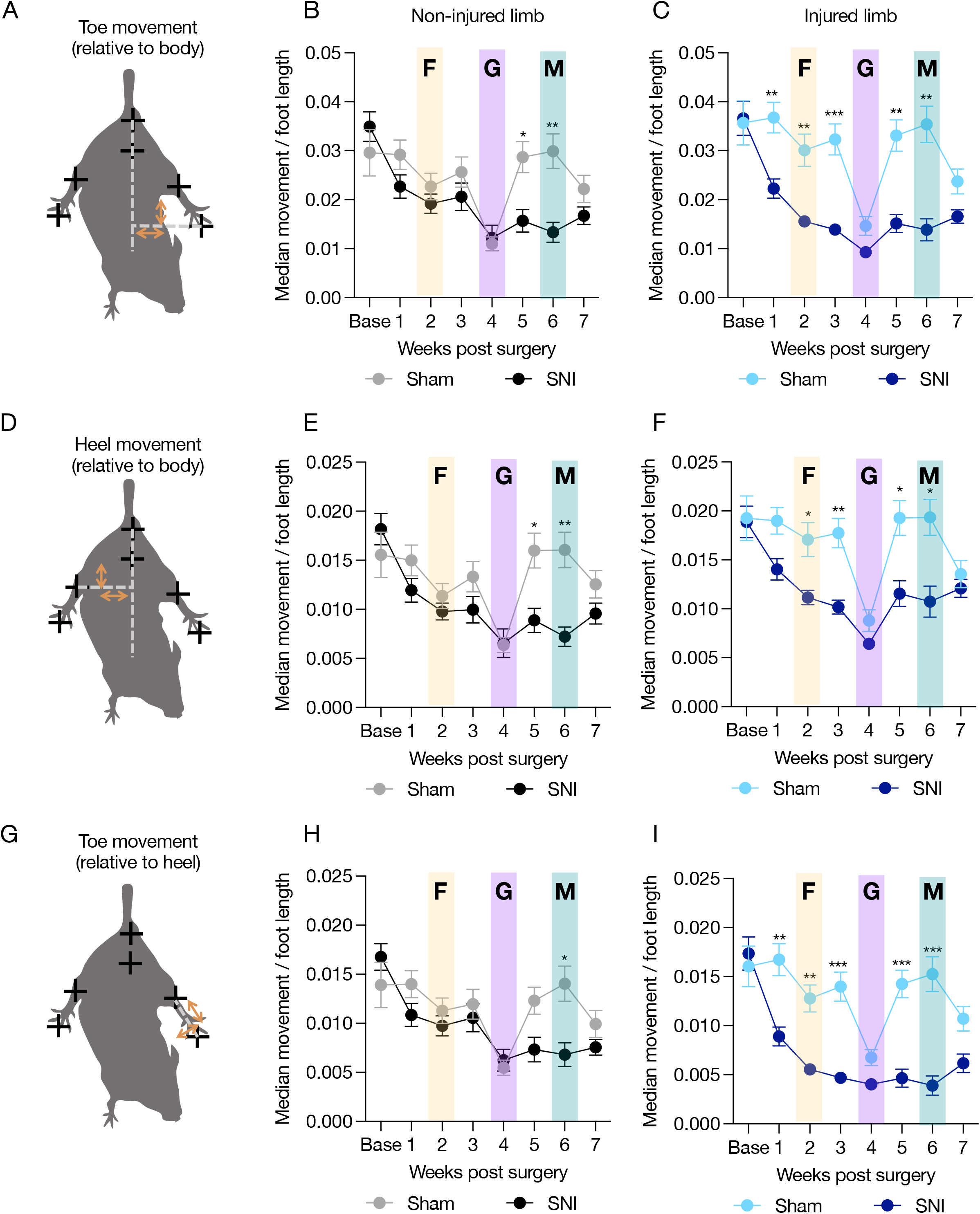
Analgesics do not reverse flail foot in injured limb. (**A**) Image shows points of comparison quantified in (**B**) and (**C**). (**B**) and (**C**) show median movement between the toe of non-injured (**B**) and injured (**C**) limbs relative to body midline. (**D**) Image shows points of comparison quantified in (**E**) and (**F**). (**E**) and (**F**) show median movement between heel of non-injured (**B**) and injured (**C**) limbs relative to midline. (**G**) Image shows points of comparison quantified in (**H**) and (**I**). (**H**) and (**I**) show median movement between the toe of non-injured **(H**) and injured (**I**) limbs relative to the heel. (**I**) shows a significant effect of SNI-induced flail foot as shown by a decrease in toe movement relative to the heel, this is independent of analgesic administration. All data expressed as mean +/−SEM; n=15-21/group; two-way ANOVA, Sidak’s post hoc (Non-injured limb Sham versus SNI; Injured limb Sham versus SNI **p<0.01 ***p<0.001, ****p<0.0001). F = Fenobam, G = Gabapentin, M= Metformin.

